# A late meta-stable code of conscious access in the absence of report

**DOI:** 10.64898/2026.01.30.702733

**Authors:** Brendan T. Hutchinson, Stanislas Dehaene, Heleen A. Slagter, Michael Pitts

## Abstract

One of the most widely recognised neural correlates of consciousness, the P3b, has been challenged in recent years as a neural signal linked to conscious processing, due to its absence in no-report paradigms. Here, we test whether, even in the absence of a P3b, a period of late metastable brain activity may provide a more general signature of conscious access to perceptual information. To this end, we leverage modern advances in electroencephalography (EEG) analyses, re-examining four datasets from no-report inattentional blindness experiments—all of which failed to find a P3b for consciously seen task irrelevant stimuli— using cross-time multivariate decoding analyses on the EEG data. We find robust temporal generalization of decoding for consciously seen stimuli across different stimulus types (shapes, faces, words), independent of report requirements and regardless of whether a P3b was evoked. This temporal generalization occurred most consistently 200-400 milliseconds post-stimulus onset, suggesting a meta-stable neural code for conscious processing in a time window typically associated with cognitive access to perceptual content. These consistent temporal generalization patterns across stimulus types indicate a potentially universal signature of conscious access.

---

One of the most robust neural signals linked to conscious processing is the P3b wave of the scalp electroencephalogram (EEG) [1, 2]. Over the past decade, this candidate “neural correlate of consciousness” (NCC) has been challenged, due in part to a series of studies by Pitts and colleagues [3–6]. These studies capitalized on “inattentional blindness”, a phenomenon in which an observer fails to notice a visual object or event that can be easily perceived if its presence is known about [7, 8]. In these studies, subjects were presented with stimuli, such as basic shapes [3], human faces [5], or letters and words [6]. Subjects were not informed of their presence and were distracted by a separate task, while EEG was used to measure their brain activity. Since stimuli were task-irrelevant and unexpected, many subjects experienced inattentional blindness to them. Yet, when subjects perceived these unexpected stimuli, but were not required to report them on a trial-by-trial basis – a so-called “no-report” condition [9] – the P3b was absent. Under these same conditions, a different electrocortical brain signal occurring earlier in time—the “visual awareness negativity” (VAN)—was instead observed [10, 11].

At the time, this series of findings was quite remarkable. It contradicted many previous studies and refuted a key assumption linking the P3b and conscious processing [10, 11] made by one of the leading neurobiological theories of consciousness, the global neuronal workspace (GNW) [12]. In each of these experiments, during post-phase questioning, many subjects retrospectively reported to have consciously perceived hundreds of stimuli, despite the complete absence of the P3b signal. This finding has since been replicated in other paradigms [13–19], with the important experimental design feature that subjects do not know in advance that a report is required about the irrelevant perceptual content. This feature is commonly labelled as “no-report” [9], but the key manipulation really concerns that of task relevancy, where stimuli that are task-relevant (whether reported or not) elicit a P3b and stimuli that are *irrelevant* to the task do not elicit a P3b, even when consciously perceived.

While these data appear to convincingly refute the P3b as a universal NCC [10, 11]), it is noteworthy that the P3b is a signal measured through univariate analyses. Univariate analyses have several well-recognized limitations, including the underlying assumption that a given signal should manifest uniformly across subjects (which contradicts the well-established heterogeneity of functional neuroanatomy [20]). Moreover, univariate analyses may not be suitable to testing more nuanced predictions of various theories of consciousness. For example, the central tenet of the GNW theory is that every conscious trial should be accompanied by a late (> 250ms) non-linear divergence between conscious and non-conscious trials (“ignition”), followed by a period of metastability lasting at least 100-200ms [21]. A particularly powerful method to overcome these limitations employs multivariate pattern analyses [22–24]. With multivariate “decoding”, a machine learning classifier is trained (to e.g., discriminate between experimental conditions) on a subset of an individual’s data and is then tested on novel data (that the classifier has not yet seen) [12, 13]. If the classifier performs above-chance at discriminating this previously unseen data, it can be inferred that information distinguishing these conditions exists within the brain, regardless of whether the spatial patterns learned by the classifier differ across subjects. In time-resolved brain data, such as EEG, multivariate decoding can be performed at every time-point to measure precisely when this information is present in the brain. Moreover, classifiers can be trained at distinct time-points and then tested across *other* time-points, an approach called “temporal generalization” of decoding [22]. With temporal generalization, it is possible to distinguish transient from “meta-stable” signals during various stages of neural processing.

Such methods have been used for a variety of purposes, including classifying the visibility of targets [24–26] (including across sense modalities [27]), distinguishing between local and global auditory novelty detection [28], and in clinical settings to monitor and distinguish states of consciousness in patients with disorders of consciousness [29]. With regards to decoding of conscious states and content, when classifiers are trained to discriminate between different classes of stimuli at each time-point in the presence versus absence of consciousness (e.g., awake versus asleep), and then tested across all time-points, a late sustained pattern of temporal generalization tends to be observed only in the conscious condition (for illustration, see Figure 1). This generalization of decoding across time resembles an off diagonal “square” (Figure 1A) or “spreading” (or “thick diagonal”) shape (Figure 1B) when consciousness is present (awake), compared with decoding only “along the diagonal” (Figure 1C) when consciousness is absent (asleep) [23].

**Figure 1.**
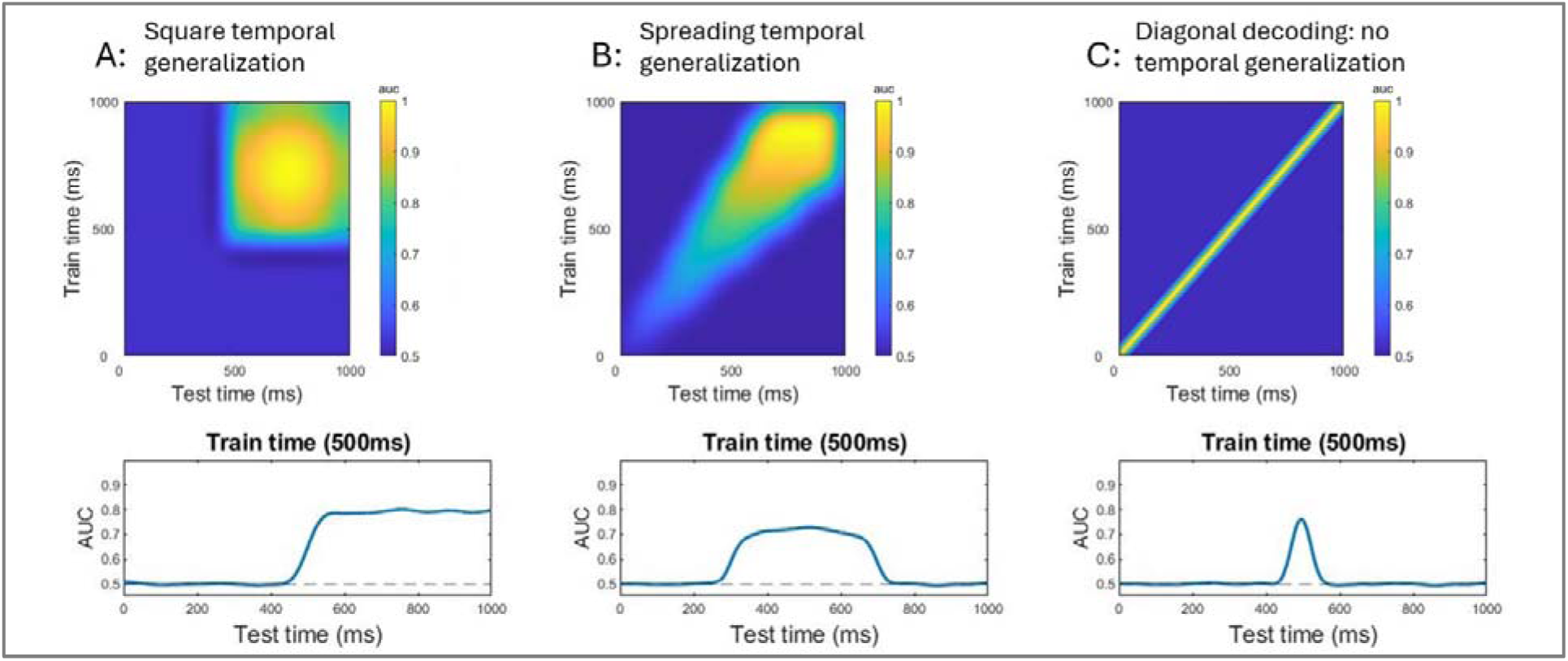
Schematic of three possible multivariate temporal generalization patterns (top row) and corresponding time series decoding of a classifier trained at 500ms (bottom row). **A:** temporal generalization emerging as a “square” shape late in time, indicating a period of meta-stability. **B:** temporal generalization increasingly “spreading” over time to create a thick diagonal, again indicating significant cross-time generalization, likely reflecting a series of meta-stable neural codes. **C:** diagonal decoding only, with little or no temporal generalization (no off-diagonal decoding), indicating a series of transient, successive neural processors.

Temporal generalization demonstrates that information earlier in time can be used to successfully decode information later in time (and vice versa), implying that the *same* (or overlapping) information is present within the brain signals across these time points. A “square” pattern of temporal generalization thus indicates that a single neural code is sustained and stabilised over time [22, 28] (Figure 1A). Empirically, when a “square” or “spreading” pattern (Figure 1B) is observed, it tends to be maximal late in time (typically > 200-300ms), suggesting it represents organized/integrated perceptual content and is not purely sensory in nature, as it occurs after initial feedforward sensory processing [22, 24]. This is also supported by the observation that late temporal generalization occurs for the content that subjects report consciously perceiving [24], even when erroneous or illusory (when not matching the veridical sensory input) [30]. Note that the simplest “square” pattern is predicted by some theoretical models when a single mental representation is consciously accessed and maintained in working memory, as observed for instance in the local-global paradigm [28]. A “spreading” or “thick diagonal” is expected, however, when conscious processing leads to the activation of a small set of successive mental representations, each lasting for a minimal duration of metastability (100-200ms). A quickly changing series of neural codes, on the other hand, should show diagonal decoding without temporal generalization (Figure 1C), for example during feedforward processing through the sensory hierarchies.

Recently, we applied multivariate decoding analyses to a no-report inattentional blindness experiment [31]. In this work, temporal generalization of decoding was robust when subjects had consciously seen the stimuli that the classifier was trained to discriminate and was minimal when these same stimuli went unseen (where decoding was mostly observed along the diagonal). Importantly, square-patterns of off-diagonal multivariate decoding were found under the same experimental conditions in which the univariate P3b was absent [31], suggesting that this late sustained pattern of temporal generalization may be present, even in the absence of a P3b. This begs the question as to whether such activity was, in fact, present in prior work where the P3b was similarly absent.

Here, we leverage data from four previous no-report inattentional blindness studies to address this question [3, 5, 6, 32]. Each used the same three-phase paradigm, where subjects were initially exposed to hundreds of unexpected suprathreshold stimuli presented at fixation, whilst performing a separate attentionally demanding task. After the first phase, subjects were questioned on whether they noticed the stimuli, and approximately half reported spontaneously noticing them (the other half being inattentionally blind). During the second phase – in which stimuli and task ran identically to the first – all subjects reported seeing the task-irrelevant stimuli, either because they had spontaneously noticed them in the first phase or because questioning had cued them to begin noticing them. In the third phase, the stimuli again remained the same, but the task was changed such that subjects were instructed to pay attention to and, in some cases, directly respond to the stimuli, rendering them *relevant* to the experimental task subjects were performing.

Examining four previous datasets offers several advantages. First, analysis of several separate datasets (totalling more than 100 subjects) greatly enhances the reliability of any given finding. Second, collectively, the *critical stimuli* for these studies were chosen from object categories that span the neural processing hierarchy, including basic shapes [3, 32], faces [5], and words/letters [6], each of which have their own dedicated processing regions and associated neurophysiological response signatures [33, 34]. A similar pattern of temporal generalization across all datasets would therefore strongly support that no unique stimulus processing architectures can explain its presence. At the same time, elements that are unique to any one study may shed light on putative heterogeneity of the late sustained activation signal that might be of theoretical interest. Third, possible confounds due to idiosyncrasies in the design or procedure of any one study can be ruled-out if the same finding is present across all datasets.

Specifically, we investigated whether multivariate EEG classifiers trained to discriminate stimuli (shapes, faces, and words/letters) from controls (random line arrays) show evidence of off-diagonal temporal generalization of decoding uniquely for stimuli that were consciously seen. The key question is whether temporal generalization is present in phase two of this three-phase paradigm, since across all studies these experimental phases were the “no-report” conditions in which irrelevant stimuli were consciously seen but did not evoke a P3b. Temporal generalization should furthermore be present in phase one for the subjects who spontaneously noticed the stimuli, but absent in phase one for subjects who were inattentionally blind to the critical stimuli. If such a pattern of results is evident, a case can be made for a late meta-stable neural signature of conscious processing that is present in the very same no-report conditions in which the P3b is completely absent [7, 10].

## Results

In each version of this three-phase no-report inattentional blindness experiment, subjects were exposed to hundreds of stimuli, each presented for 300ms and interspersed with random line arrays for 533-800ms (with slight variability between studies) (see Figure 2). Critical stimuli in the first study (datasets one and two [3, 32]) were square patterns formed from line arrays (Figure 2B left). In study two (dataset three [5]), critical stimuli were human faces generated from random lines (Figure 2B middle). In study three (dataset four [6]), words and consonant strings were formed from random lines (Figure 2B right). EEG was recorded during the presentation of these stimuli across three experimental phases in which *awareness* and *task relevancy* of the critical stimuli were experimentally manipulated. In phase one, stimuli were either spontaneously seen (*aware*) or went unnoticed (*inattentionally blind*), but there were no a priori report requirements for the stimuli, rendering them *task-irrelevant*. In phase two, stimuli were always seen but remained task-irrelevant, hence no-report requirements persisted in this phase. In phase three, stimuli were consciously seen and *task-relevant* (Figure 2A).

**Figure 2.**
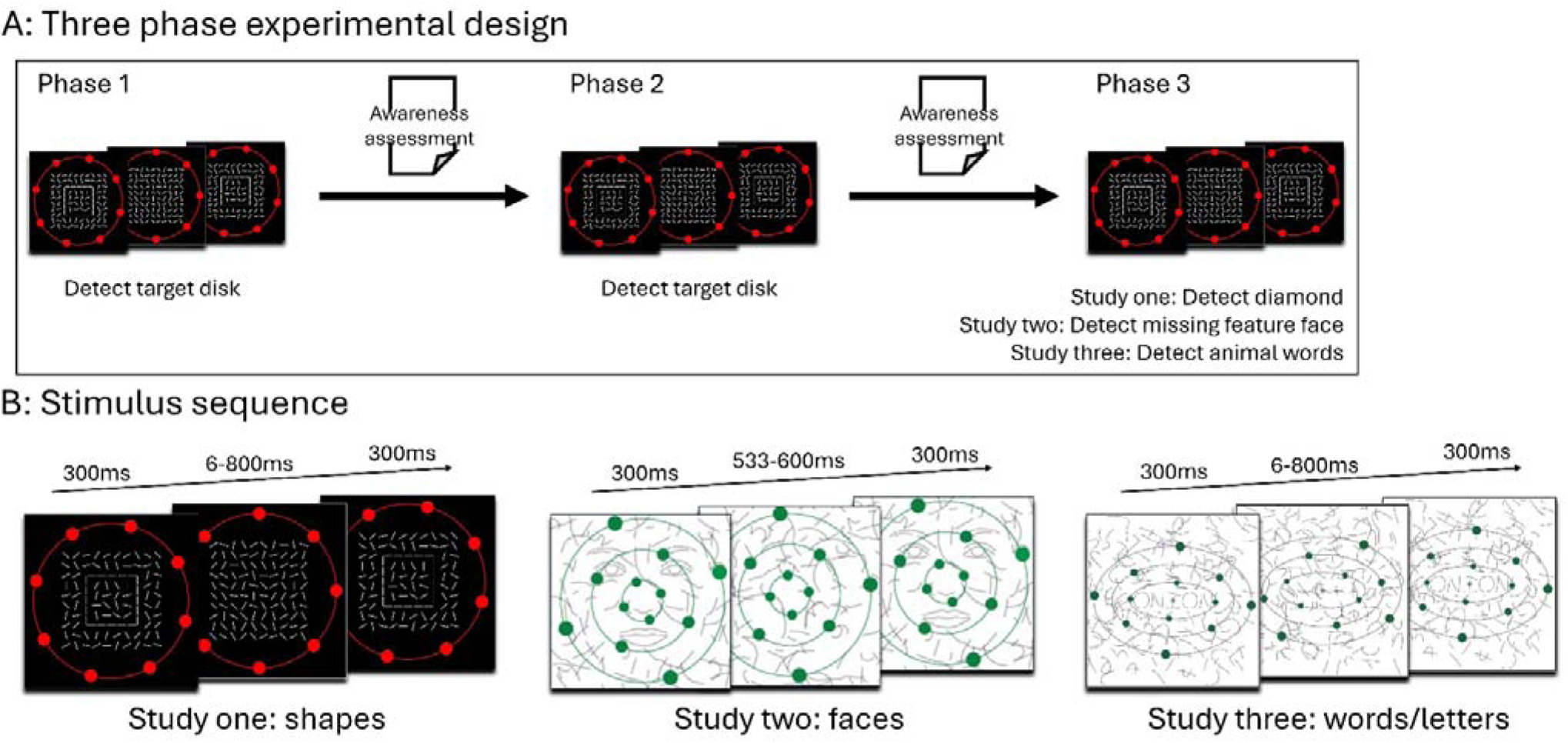
Different versions of the no-report inattentional blindness experiment used in studies 1-3 (datasets 1-4). **A)** Overview of three-phase design. Subjects perform a distractor task in which they have to respond to an infrequent target disk in phases one and two. A task switch occurs in phase three where subjects instead have to respond to infrequent target shapes (study one), faces (study two), or words (study three) presented centrally as a part of the line arrays. An assessment of subjects’ awareness of the centrally presented unexpected stimuli occurs in between phase one and two, and again between phase two and three. Study one shown here for illustration purposes. **B)** Stimulus sequence used in all experiment phases for studies one to three. The presentation of unexpected stimuli (300ms) is interspersed with a screen of random line arrays for a variable duration (study one/three: 600-800ms, study two: 533-600ms). This sequence is repeated until the end of a block (∼1-2 mins). Distractor stimuli (study one: red ring of disks, study two/three: green rings of disks) continue to rotate, either in a jittered manner with every central stimulus refresh (study one) or with continous smooth motion (study two/three), until the end of a block.

We first sought to assess performance of classifiers trained to discriminate critical stimuli from their controls (random lines) in phase three, when subjects were consciously attending to and actively discriminating between stimuli. It has been shown in every iteration of this no-report inattentional blindness experiment that task-relevant stimuli in phase three evoke a P3b (typically onsetting from about 300-350ms) [3–6, 32] (supplementary Figure 1 shows complementary VAN/P3b data from [38]). Performance in this phase therefore cannot demonstrate a dissociation between P3b and temporal generalization of decoding, but it can serve as an initial validation step by indicating the strength and shape of decoding for consciously seen task-relevant stimuli that elicit a P3b. Several previous studies using only report-based methods have found a late square-shaped pattern of temporal generalization for consciously perceived stimuli [24–26, 35, 36]. Beyond being important for validation and replication purposes, we reasoned that analyses of phase three data would provide a plausible ceiling to decoding performance and hence aid with interpreting classifier performance under the more stringent and theoretically interesting no-report conditions of phase two and phase one.

### Consciously seen stimuli in report conditions produce strong temporal generalization

As Figure 3 shows, there was robust temporal generalization in phase three of each study. The temporal generalization here can be characterized as a series of successive “square” patterns, that began from ∼ 150ms and persisted through to the end of the epoch in all studies (Figure 3). For study one (shapes), although performance peaked around 300ms in terms of sheer magnitude (peak AUC = 80.14%, *p* < .001), off-diagonal decoding improved with time (in terms of temporal spread), as shown in Figure 3 (middle column), which displays off-diagonal decoding as a function of training time (200-800ms, in steps of 100ms). While the generalisability of the classifier spanned from 152 to 292ms when inspecting classifiers trained at 200ms (*p* < .001), classifiers generalized for close to half a second (540- 956ms) when inspecting classifiers trained at later times of e.g., 700ms (*p* < .001) (Figure 3A). For all studies, temporal spreading was extensive at time windows around the P3b: classifiers trained at 400ms generalised from 320-536ms for shapes (peak AUC = 77.41%, *p* < .001), 148-692ms for faces (peak AUC = 85.32%, *p* < .001) and from 108-544ms for words/letters (peak AUC = 83.17%, *p* < .001) (Figure 3B, C). Posterior sensors contributed most to classification performance in time windows from 200-500ms: bilaterally for shapes and words/letters and lateralised over the right hemisphere over temporoparietal regions for faces. For shapes and faces, maximal performance spread to central-parietal locations at later time points (∼500-600ms) (refer to scalp maps, Figure 3).

**Figure 3.**
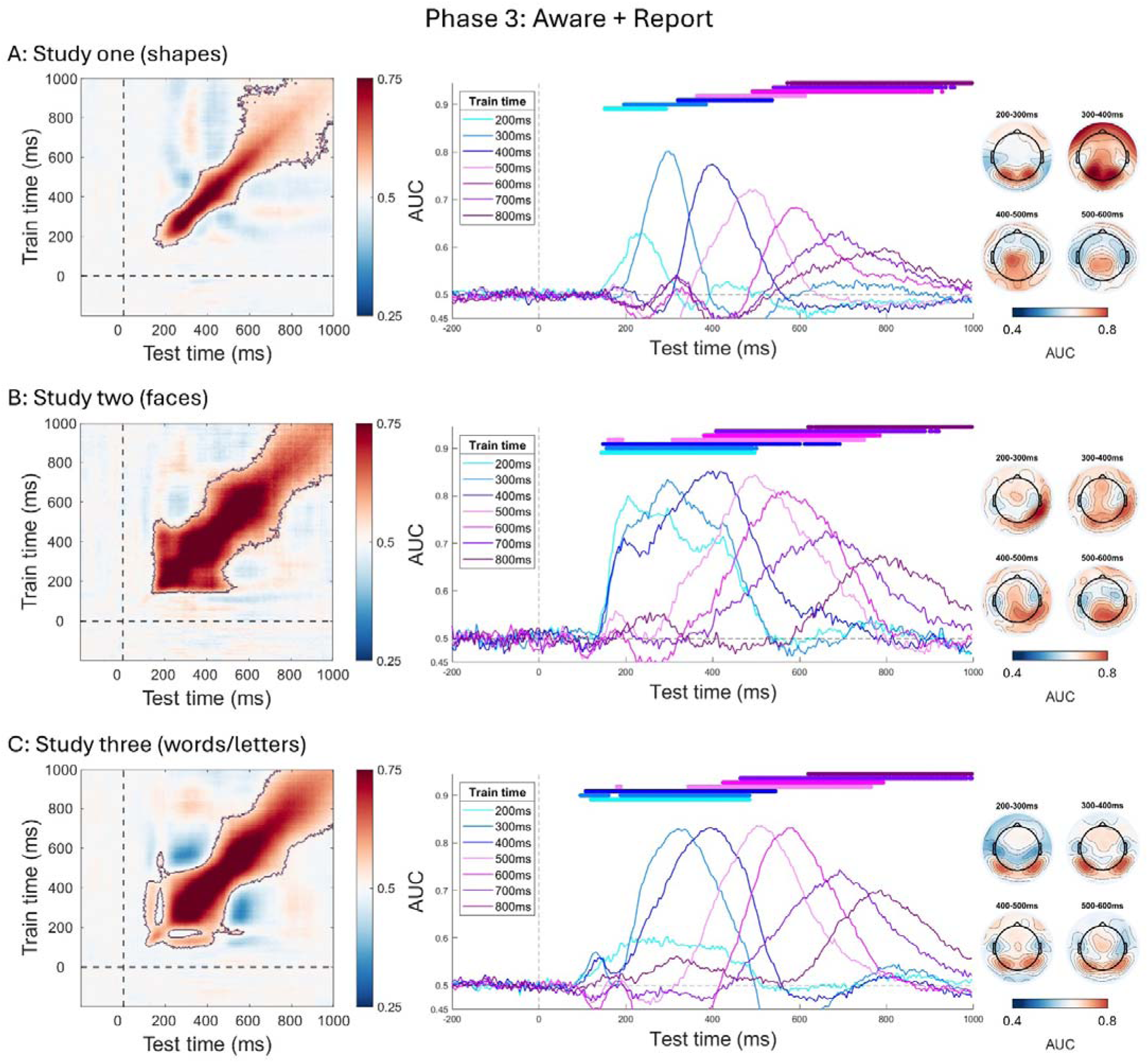
Multivariate decoding results from phase three of each study (**A:** shapes, **B:** faces, **C:** words/letters). Heatmaps (left) show temporal generalization of decoding, where classifier training time is represented along the y axis and testing (generalization) time is along the x axis. Classification performance (AUC) is indicated by colour intensity of the heatmap, where statistically significant temporal generalization is outlined. Time series plots (middle) show performance of classifiers trained at distinct time points from 200-800ms in 100ms intervals. Colour transitions here indicate different training times and areas of statistically significant classification performance are indicated by (colour coded) bars above the time series. Scalp maps (right) show classification performance based on a searchlight analysis, where classifiers are trained sensor-by-sensor in distinct time windows of 100ms length from 200-600ms.

### Temporal generalization of seen stimuli in no-report conditions

After confirming robust temporal generalization of decoding for consciously seen stimuli under report conditions (when stimuli evoke a P3b), we next sought to address the more critical question of whether consciously seen stimuli under no-report conditions (that do *not* evoke a P3b) also show evidence of temporal generalization of decoding. Strikingly, when we trained classifiers to decode critical stimuli from their controls in phase two, while there was a (to-be-expected) decrease in absolute magnitude of performance compared with phase three, the initial pattern of generalization under no-report conditions resembled that under report requirements, particularly from ∼200-600ms (Figure 4).

**Figure 4.**
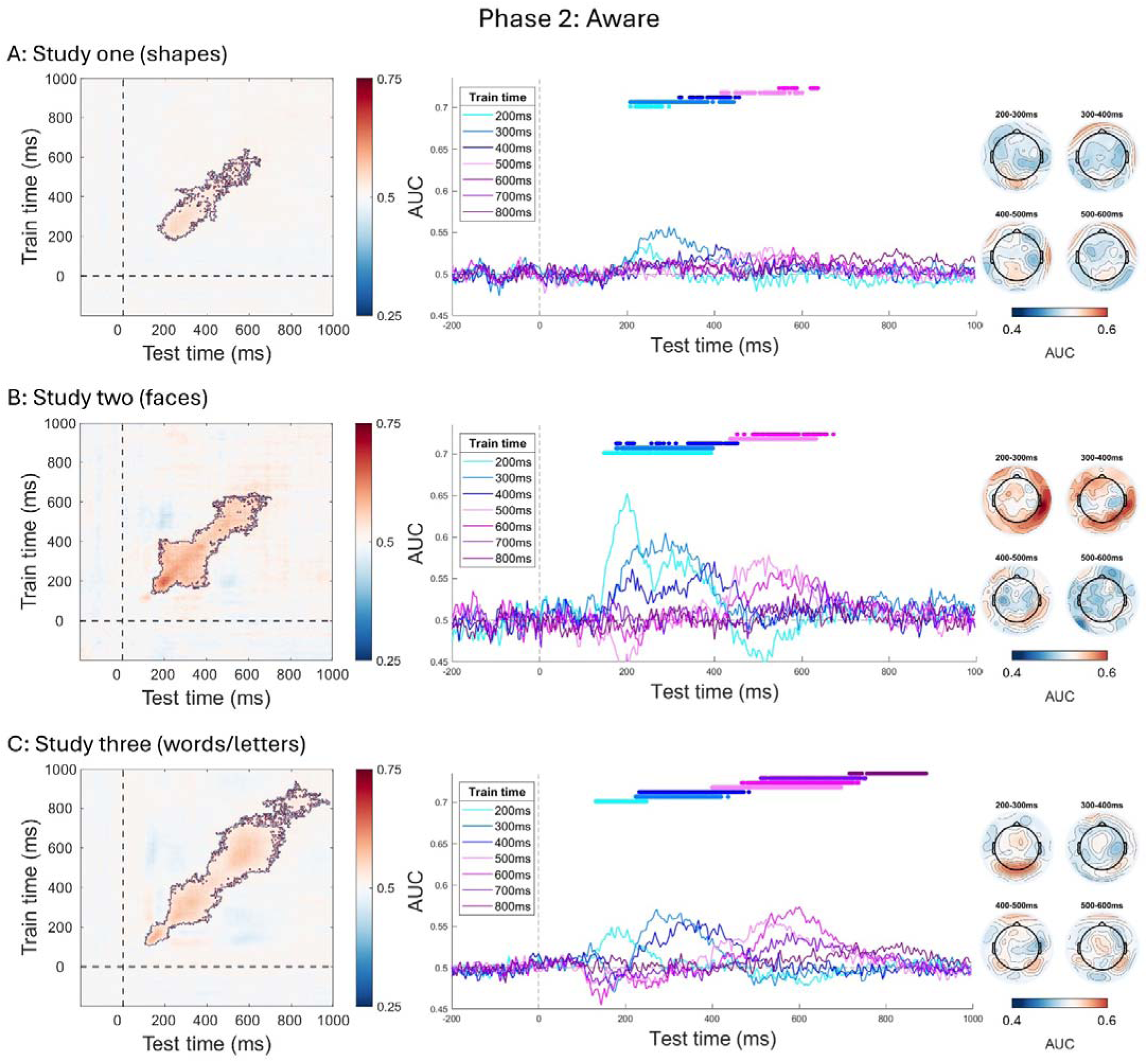
Multivariate decoding results from phase two of each study (**A:** shapes, **B:** faces, **C:** words/letters). Heatmaps (left) show temporal generalization of decoding, where classifier training time is represented along the y axis and testing (generalization) time is along the x axis. Classification performance (AUC) is indicated by colour intensity of the heatmap, where statistically significant temporal generalization is outlined. Time series plots (middle) show performance of classifiers trained at distinct time points from 200-800ms in 100ms intervals. Colour transitions here indicate different training times and areas of statistically significant classification performance are indicated by (colour coded) bars above the time series. Scalp maps (right) show classification performance based on a searchlight analysis, where classifiers are trained sensor-by-sensor in distinct time windows of 100ms length from 200-600ms.

Importantly, in each study, there was clear evidence of temporal generalization of decoding: off-diagonal decoding peaked most consistently at training time points around 300ms in temporal spread but was nevertheless present when classifiers were trained from 200-600ms. For example, classifiers trained at 600ms for study three (words/letters) showed significant decoding from 480-724ms (peak AUC 57.40%, *p* < .001), although off-diagonal decoding was observed for training times even up to 800ms in this dataset (Figure 4C). Notably, off-diagonal decoding was robust across all studies in the 300-400ms time window when the P3b is typically evoked: training times of 300ms for study one (shapes), 208- 444ms, peak AUC = 55.69%, *p* = .003 (Figure 4A, middle column), and study two (faces), 176-396ms, peak AUC = 60.52%, *p* < .001 (Figure 4B, middle column); and training times of 400ms for study three (words/letters), 232-472ms, peak AUC = 55.78%, *p* < .001 (Figure 4C, middle column). Like the report conditions of phase three, posterior sensors contributed most to temporal generalization under the no-report conditions of phase two—predominantly right hemispheric temporoparietal sensors for faces and more central occipital sensors for study one (shapes) and three (words/letters) (see scalp maps of Figure 4).

Next, we turned to the data from phase one, which we expected to present differently from phase two and phase three since, in each study, about half of the subjects were inattentionally blind to the critical stimuli in this phase. This means that analyses of the phase one data will invariably be lower powered compared with phase two and phase three, as the splitting of subjects into two groups effectively halves analytic power. More importantly, because awareness is assessed in a phase-wise fashion in this paradigm—and the moment in which spontaneous noticer subjects may first notice the critical stimuli varies (i.e., during earlier or later blocks of trials within this phase)—the sheer number of “aware” trials in phase one is likely considerably lower than any other phase for this group of subjects. In effect, we reasoned that any positive effects for seen stimuli in phase one *underestimates* the true extent of temporal generalization of decoding for seen stimuli due to contamination of trials labelled as “aware” with those that were unseen (a well-recognised limitation of the no-report inattentional blindness paradigm [7]).

Despite these constraints, temporal generalization of decoding was still observed in phase one for those subjects who spontaneously noticed the unexpected critical stimuli, where classifiers trained from ∼200-400ms showed significant decoding (Figure 5), again with peak performance most consistent for classifiers trained at 300ms, although late classifiers also showed off-diagonal decoding at training times of ∼500-700ms for study three (words/letters) (Figure 5C). Once again, posterior sensors contributed most to classification performance in these time windows.

**Figure 5.**
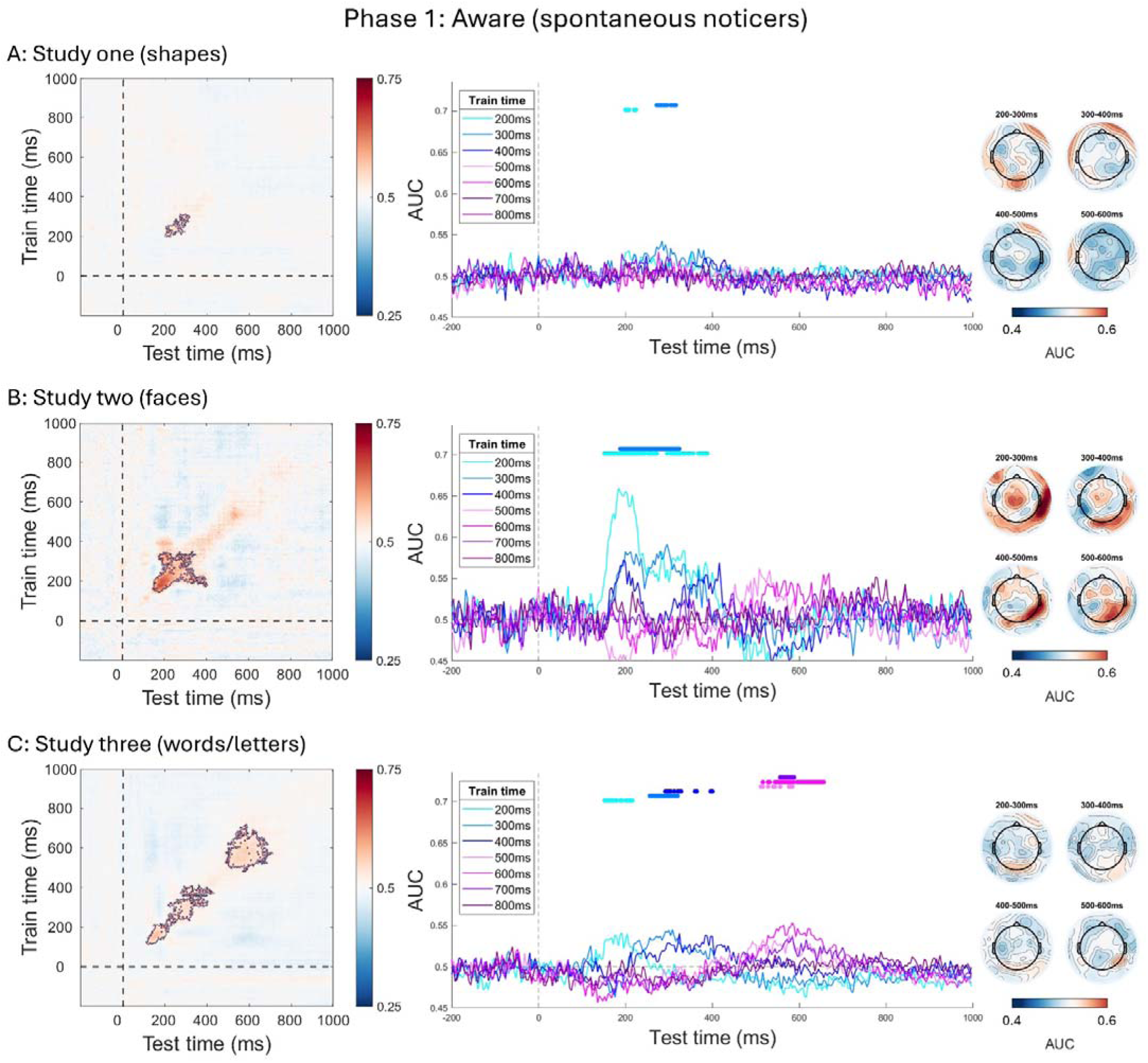
Multivariate decoding results from phase one (aware subjects) of each study (**A:** shapes, **B:** faces, **C:** words/letters). Heatmaps (left) show temporal generalization of decoding, where classifier training time is represented along the y axis and testing (generalization) time is along the x axis. Classification performance (AUC) is indicated by colour intensity of the heatmap, where statistically significant temporal generalization is outlined. Time series plots (middle) show performance of classifiers trained at distinct time points from 200-800ms in 100ms intervals. Colour transitions here indicate different training times and areas of statistically significant classification performance are indicated by (colour coded) bars above the time series. Scalp maps (right) show classification performance based on a searchlight analysis, where classifiers are trained sensor-by-sensor in distinct time windows of 100ms length from 200-600ms.

### Minimal temporal generalization of unseen stimuli during inattentional blindness

After demonstrating consistent temporal generalization of decoding for all phases in which subjects were aware of the critical stimuli, including under no-report conditions where stimuli did not evoke a P3b, we asked whether any temporal generalization would emerge when stimuli were unseen due to inattention. In the inattentionally blind group of subjects in phase one, there were no significant clusters of multivariate decoding observed for study one (shapes) or two (faces), nor did any sensors seem to contribute to classification performance (Figure 6). For study three (words/letters), there was a time window of significant decoding observed from ∼112-344ms. Classifiers trained at 200ms and 300ms showed some sparse off diagonal decoding for about 50ms (156-160ms, 168-176ms, 192ms, 200-208ms, *p* = .02) and 120ms (244-260ms, 272-364ms, *p* = .01), respectively. Hence, during inattentional blindness, two out of the three studies (shapes, faces) showed no significant decoding, while one (words/letters) showed some significant decoding, but with much reduced temporal generalization compared with phase two (where all subjects were aware of the critical stimuli) and only relatively early, before the P3b time window.

**Figure 6.**
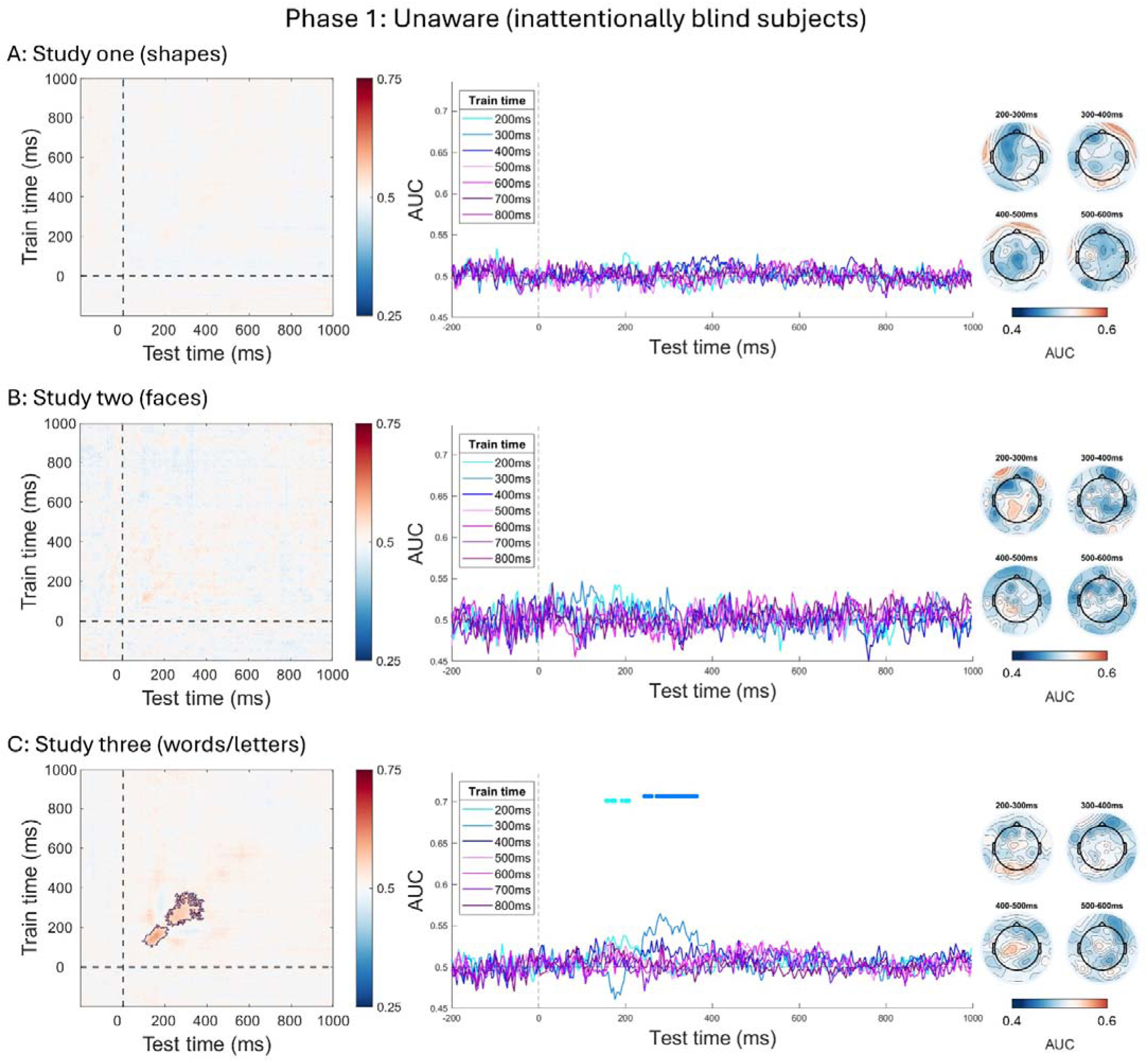
Multivariate decoding results from phase one (inattentionally blind subjects) of each study (**A:** shapes, **B:** faces, **C:** words/letters). Heatmaps (left) show temporal generalization of decoding, where classifier training time is represented along the y axis and testing (generalization) time is along the x axis. Classification performance (AUC) is indicated by colour intensity of the heatmap, where statistically significant temporal generalization is outlined. Time series plots (middle) show performance of classifiers trained at distinct time points from 200-800ms in 100ms intervals. Colour transitions here indicate different training times and areas of statistically significant classification performance are indicated by (colour coded) bars above the time series. Scalp maps (right) show classification performance based on a searchlight analysis, where classifiers are trained sensor-by-sensor in distinct time windows of 100ms length from 200-600ms.

To ensure temporal generalisation in aware subjects of phase one could not be attributed to any difference in power between groups (due to differing sample sizes in the aware versus inattentionally blind groups), we bootstrapped a sub-sample of the aware subjects (study one: *N* = 32, study three: *N* = 28) to match the number of inattentionally blind subjects (study one: *N* = 29, study three: *N* = 18) (we did not perform this analysis for study two as sample sizes were identical in both groups for this study). We performed resampling 100 times, selecting a new random sub sample on each iteration. Results are presented in supplementary figure 3 and show the same pattern of temporal generalization in aware subjects, even when the sample size was matched with that of the inattentionally blind group of subjects.

### Cross-phase generalization: decoding of seen stimuli in report conditions generalizes to no-report conditions

In addition to the above analyses, which were aimed at classifying stimuli within each phase, the three-phase experimental design also provides an opportunity to decode across phases. We considered such analyses useful to test whether decoding of stimuli in report conditions generalizes to no-report conditions, which could provide corroborative evidence that temporal generalization is uniquely associated with conscious perception.

We trained classifiers to discriminate stimuli (critical stimulus versus control) in phase three and then tested these same classifiers on both phase two and phase one (separately for spontaneous noticers and inattentionally blind subjects), thus asking whether the pattern of neural activity when stimuli were attended and task-relevant generalizes to the no-report conditions when stimuli were task-irrelevant (for cross-phase generalization from phase two to phase one, refer to supplementary figure 2). Our rationale was that, while it remains possible that activity associated with report and working memory may dominate classification performance in phase three (and hence not generalize to phase two or one), neural activity in response to stimuli in this phase nevertheless showed the highest signal-to-noise ratio (as evidenced by classification performance in phase three), and thus training on this phase may provide a good opportunity to assess how, if any, awareness-related activity may generalize to other experimental phases.

Figure 7 shows results for each study. In all, temporal generalization of decoding was observed for classifiers trained on phase three and tested on phase two, most consistently from ∼200-400ms but with some additional, slightly asymmetrical, activity at later time points for study one (shapes) and three (words/letters) (Figure 7A and C, left column). The same was true, but to a lesser extent, for classifiers trained on phase three and tested on phase one in the spontaneous noticers (Figure 6, middle column). Largely no cross-phase generalization was evident from phase three to phase one for subjects who were inattentionally blind in phase one, except for a brief bout of decoding in study three (words/letters) at testing time points of about ∼180ms. Interestingly, such decoding was also asymmetrical: classifiers trained on phase three could be trained at any time between ∼180ms and 400ms and yet generalize to this brief period of non-conscious phase one data only around ∼180ms. This finding is consistent with the hypothesis that unseen stimuli can cause brief activations, which become sustained over time only when the threshold for conscious perception is crossed [28].

**Figure 7.**
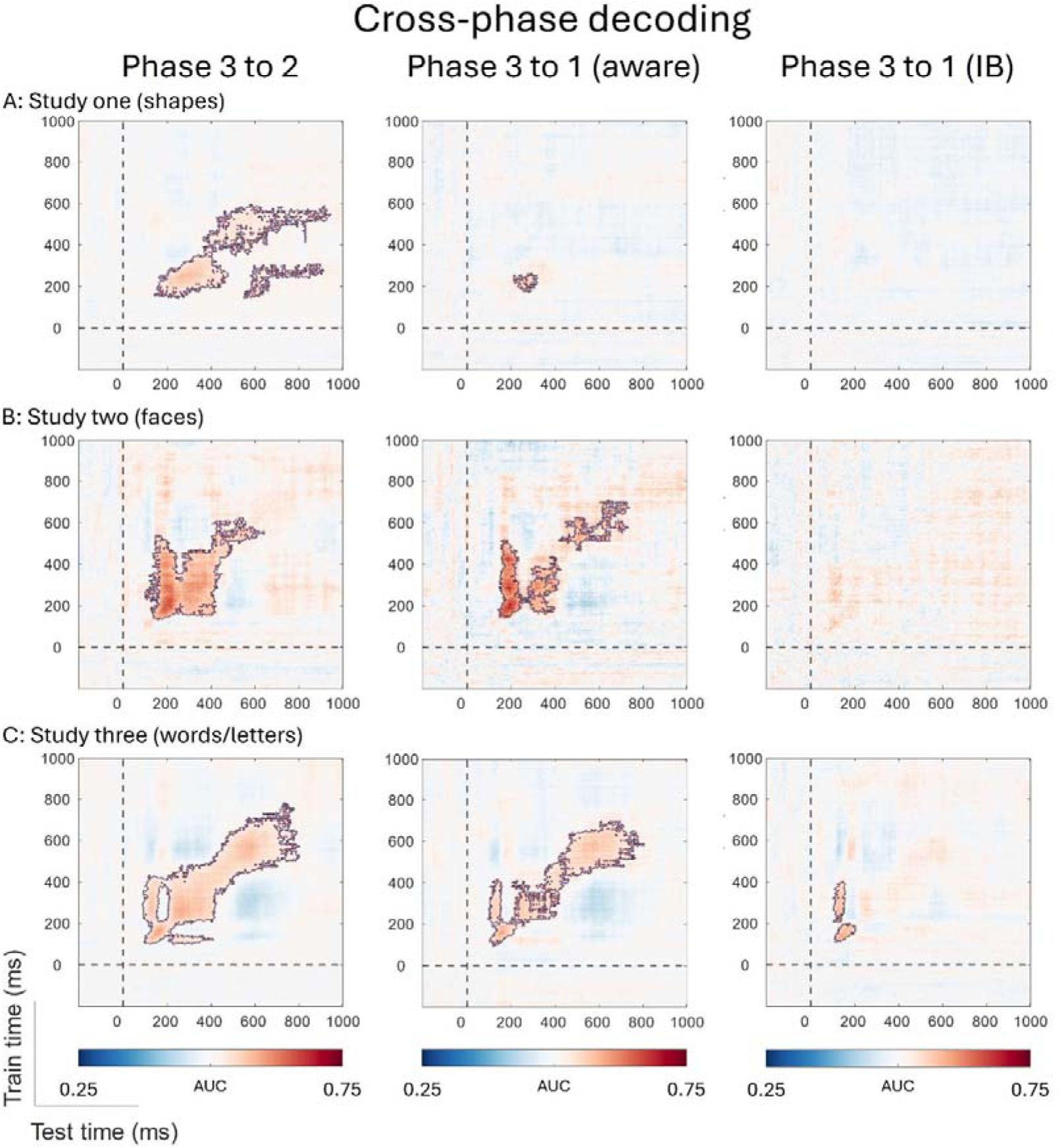
Cross-phase decoding results, where classifiers were trained on the report condition, phase three, and tested on the no-report conditions, phase two (left) and phase one, separately for spontaneous noticers (‘aware’, middle) and inattentionally blind subjects (‘IB’, right), for each study (**A:** shapes, **B:** faces, **C:** words/letters). Classifier training time is represented along the y axis and testing (generalization) time is along the x axis. Classification performance (AUC) is indicated by colour intensity of each heatmap, where statistically significant decoding is outlined.

To complement the above, we also examined the difference in temporal generalisation between phase one and phase two in the inattentionally blind group of subjects (we could not use aware subjects for this analysis because they do not have their own unseen control condition), thereby directly contrasting the patterns of temporal generalisation for the same stimuli when they were seen (phase two) versus unseen (phase one) in the same group of subjects. Results of these analyses are presented in supplementary figure 4 and show bouts of decoding that was significantly stronger in phase two (when stimuli were seen) compared to phase one (when stimuli were unseen) for two out of three studies (study one and study two).

## Discussion

Consciously seen stimuli produced robust patterns of temporal generalization of decoding independent of report requirements. These patterns were evident across four datasets, all of which used variants of a three-phase no-report inattentional blindness experiment [3, 5, 6, 32]. Temporal generalization of decoding was strongest and longest lasting in time in the phase in which stimuli concomitantly evoked a P3b—when they were task-relevant and had to be actively discriminated—yet generalization was still present during the experimental phases where the P3b was absent (under no-report conditions) and extended into—and sometimes beyond—the time window in which the P3b typically occurs. This same result was largely absent when subjects were unaware of the critical stimuli due to inattentional blindness. This late meta-stable pattern generalized from the report condition (phase three) to the no-report conditions (phase two and one) only when subjects were aware of the stimuli (barring some minor asymmetrical off-diagonal decoding for words/letters, see Figure 7). We interpret these findings to indicate that a meta-stable code of stimulus processing is activated most prominently from about 200ms to 400ms post-stimulus-onset, independent of stimulus type and task/report conditions, only when that stimulus is consciously accessed. In some cases, temporal generalization clearly extended beyond this time (e.g., up to 800ms for words/letters in no-report conditions), suggesting possible variation in the temporal extent of this effect across stimulus types.

While debate persists on the precise timing at which genuine conscious processing can be read-out from human brain recordings [10], the 200-400ms post-stimulus time window has previously been considered critical in terms of signifying the change from pre- conscious to conscious processing [37]. It is somewhat earlier than the late-stage cortical ignition initially proposed by the GNW model [38] yet is later than speculated by some sensory theorists who link consciousness with early stage (i.e., 100-200ms) recurrent processing in sensory regions [39]. This intermediate time window is therefore somewhat complementary to both positions, in that it reflects a middle ground between these two extremes. Moreover, the timing and relative scalp topography of decoding (determined here via searchlight analyses) coincides well with the “visual awareness negativity” (VAN), an electrocortical brain signal that was previously observed in each dataset we analysed here, in a time window closely mapping to this 200-400ms time window (VAN timing in study one: 300-340ms [3], study two: 265-305ms [5], study three: 320-380ms [6]). A more anterior N2 (coupled with a late bilateral occipital positivity) is also often associated with conscious processing in a similar time window within other paradigms [e.g., 16, 40, 41] and similarly maps well to this time window. One plausible interpretation of our findings is therefore that they reflect the multivariate counterpart to the VAN (or the frontocentral N2), since temporal generalization of decoding was observed in all phases where the VAN (or N2) was previously observed (at approximately the same time window). However, the duration of the VAN/N2 is typically much shorter than the meta-stable patterns of decoding observed here and, importantly, generalization clearly extended beyond this time window in these same experimental conditions. Therefore, this temporal generalisation signature is considerably more sensitive than the univariate VAN/N2.

A larger question is whether these findings are more in line with sensory models of consciousness [39, 42] or cognitive models such as the GNW [1–2, 12, 38] or higher order theories [43]. According to the GNW model, late temporal generalization of decoding reflects cortical ignition and the brain-wide broadcasting of perceptual content. The very nature of temporal generalization implies that the underlying signal is a robust neural representation that persists across time – one that, according to GNW, is sustained by a single mechanism (e.g., attention and/or working memory) – and which can be read out from scalp recordings after initial feedforward sensory processing has taken place [22]. These observations fit with temporal generalization as an ideal marker of conscious processing, and may suggest “global broadcasting”, in line with the workspace model of consciousness [12, 23, 44].

That said, if the meta-stable pattern was truly “globally” broadcast, one might expect to see a broader spatial distribution of decodability than what was observed here, since we found a mainly (albeit not exclusively) posterior focus of decoding across all three paradigms in the no-report conditions. Temporal generalization also appeared to change substantially over time, as we observed distinct “waves” of off-diagonal decoding, rather than one extended process, which conceivably reflects changing scalp topographies (and possibly meta-stable codes) over time. Along the same lines, some sensory theories of consciousness, such as the integrated information theory (IIT) also predict temporal generalization of decoding, albeit only in posterior cortical regions and only persisting for the duration of the conscious experience [21, 45]. Future research using more spatially precise methods of decoding (e.g., with fMRI or intracranial EEG data) and including manipulations of stimulus duration coupled with manipulations of perceptual awareness will be necessary to fully evaluate whether these meta-stable patterns of decoding are more consistent with specific cognitive or sensory theories of consciousness [21].

A recent fMRI study found that, when analysed with univariate statistics, differences in activation of the prefrontal cortex between seen and unseen conditions were present in the report condition but were completely absent in the no-report condition [46]. However, multivariate analyses (decoding) of this same contrast (seen versus unseen) revealed statistically significant decoding in the prefrontal cortex, including in the no-report condition [see also 47]. These results appear to parallel the pattern of findings here: a *multivariate* signature of conscious awareness occurred in the absence of the *univariate* signal (i.e., the P3b). Again, however, while our current results conclusively identified a meta-stable period of neural activity linked with conscious perception, given the spatial limitations of scalp-recorded EEG, we cannot make definitive claims regarding the location or underlying neural generators of these patterns. Moreover, it is not entirely clear what features of the EEG signal our classifiers used to discriminate between stimulus conditions, and so it is difficult to give any precise description of the neurophysiological mechanism underlying these temporal generalisation patterns.

Our findings hold significant implications for broader research on the neural correlates of consciousness, irrespective of whether seen through the perspective of the GNW model, sensory theories, or a more theoretically neutral lens. If GNW’s account holds, then our findings, which suggest meta-stability of perceptual content beyond the time-window of the VAN, strongly counter the conclusions drawn from the past decade of literature that has relied on the absence of a P3b for seen stimuli in no-report paradigms to refute the workspace model [10]. Such a finding has been made in various studies, ranging from the inattentional blindness experiments reported on here, to masking [13] and more recently, the attentional blink [14]. Several authors have concluded that this absence of the P3b refutes the GNW outright [10, 48]. However, it remains possible that the GNW predictions of non-linear ignition and global broadcasting are accurate but that these events are simply not always signalled by a P3b (i.e., “ignition” without a P3b) [16]. Furthermore, it is important to note that, even though it has been experimentally separated from consciousness, the P3b nevertheless remains useful in a clinical context in identifying patients with residual consciousness, as it seems to index a subset of conditions that nevertheless indicate conscious processing, for instance conscious access to working memory [49].

If GNW’s account holds, an important future consideration will be addressing how these findings might fit within the GNW model. Recently, another group of researchers made a similar observation, where temporal generalization of decoding was reported for task-irrelevant sounds that were consciously perceived, yet with no concomitant P3b [50]. The authors argued that their findings may suggest the workspace can be active with “no specific agenda”, referring to this subcomponent of the workspace as a “global playground”. In other words, a brief bout of meta-stability during intermediate time ranges might always be necessary for conscious access, while stronger and more extended meta-stable patterns are optional and depend heavily on how the perceptual information is being used to meet current task-demands. It is feasible that our findings reflect a similar process, although more work is needed here before firm conclusions can be drawn.

From the perspective of a sensory theorist (e.g., recurrent processing theory or IIT), the current results might reflect a stage of local recurrent processing or perceptual integration necessary for phenomenal consciousness, whether or not such information is ever accessed by cognitive systems [39, 42, 51]. Indeed, recent data from three separate intracranial EEG studies indicated robust temporal generalization of decoding in posterior visual areas that matched the duration of task-irrelevant stimuli [21, 52, 53]. Sensory theories that posit the existence of phenomenal consciousness without access, however, would need to explain how subjects in the current set of experiments were so easily able to retrospectively report what they saw during the post-phase awareness assessments if the perceptual information (from the critical stimuli) was never accessed.

If one adopts a more theoretically neutral lens, then our findings demonstrate at minimum a late multivariate signature of conscious perception under no-report conditions. Workspace theorists previously speculated that such patterns of multivariate decoding may uniquely reflect conscious processing [23], but to our knowledge, the results reported here are some of the first to directly demonstrate temporal generalization of decoding under no-report conditions that is present when subjects are aware of the stimuli and largely absent when they are not [see also 31, 50, 54]. The fact that these patterns of temporal generalization were so similar across different iterations of the three-phase inattentional blindness experiment, despite changes in the critical stimuli, suggests some consistency to the multivariate decodability of conscious content more broadly. Indeed, it is remarkable that a decoder trained on phase three, where the stimuli were task-relevant and generated a P3b, could generalize to phase two, a no-report condition where the same stimuli were task-irrelevant and did not evoke a P3b. Such generalization of decoding demonstrates that, in spite of the large changes in scalp ERPs, a common metastable neural code was activated in both conditions. Intriguingly, in all three studies, the results from phase two suggested two or more successive stages of meta-stable neural codes. If such a pattern is verified in future studies, using different manipulations of awareness, this could lend support to a recent multi-stage proposal of conscious processing that could potentially unify certain aspects of sensory and cognitive theories of consciousness [55].

Still, the results varied somewhat across the three experiments, which used distinct stimuli (shapes, faces, and words/letters). Patterns of decoding related to more complex stimuli (faces and words/letters) undeniably emerged as more robust patterns than basic shapes, which instead showed a tighter funnelling and less spreading late in time. It will be of interest in future studies to determine whether these precise patterns and their differences across stimulus types are meaningful. One plausible hypothesis is that more complex stimuli will inevitably show greater temporal generalization because these stimuli are perceptually more easily distinguishable from control stimuli and require more extensive neurophysiological processing at higher stages of the visual cortical hierarchy [33, 34].

Potential methodological reasons will also need to be ruled out. For example, both datasets we examined that used more complex stimuli (faces and words/letters) also employed higher density EEG scalp recordings (96 electrodes), which meant the classifier used higher resolution electrocortical data to discriminate between conditions compared with the datasets with simpler stimuli (shapes), which were recorded from fewer electrodes (64 electrodes). One relatively simple follow-up could be to assess whether the patterns of decoding we observed here for shapes can be replicated with higher resolution scalp recordings.

Could off-diagonal decoding have a simpler explanation, one that would occur even on non-conscious trials? One possibility is that temporal generalization here reflects variability in the latency of evoked responses. For example, ERP components – particularly those occurring later in time – tend to be more temporally “smeared” due to, e.g., intertrial variability in component latencies [57]. The same could be true for multivariate processes. Under this logic, any jitter in the evoked response may be captured in the multivariate decoding response. We ultimately find this interpretation unlikely, since off-diagonal decoding here also occurred early in time and extended for too long in time to be easily explained by inter-trial variability of evoked potentials. For example, off-diagonal decoding of faces under no-report conditions (study two) clearly showed that training a classifier at 200ms resulted in temporal generalization from about 180ms to 400ms (see Figure 4B). ERP components that cover such a wide and varied latency range are typically endogenous and generally occur later in time, such as the P3b, N400, or the late positive potential (LPP).

Indeed, the fact that temporal generalization occurred in the time window of the P3b—in the absence of a P3b—suggests that the more plausible interpretation is that the off-diagonal decoding observed here captured the neural representation of the perceptual content itself, in this case a face.

An important question that remains is whether temporal generalization can reliably occur for unseen stimuli. No observable multivariate decoding was observed in unseen conditions for three out of four datasets we analysed (see Figure 5). However, significant decoding for unseen stimuli in phase one was observed in one of the datasets (from ∼112- 344ms), where words and consonant strings were used as the critical stimuli. We similarly found significant decoding of unseen words in a previous no-report inattentional blindness experiment [31, see also 58]. In both cases, decoding was mostly (albeit not exclusively) constrained to diagonal (same time) classifiers, which is thought to reflect more serialised and modular processes that do not require stabilization [22]. That said, since diagonal decoding was observed (for unseen stimuli) for two out of five datasets (including [31]), one could argue that meta-stability of a representation has not been wholly demonstrated, since *any* form of decodability at all may differentiate seen from unseen stimuli. It will therefore be important in future work to more thoroughly establish whether unseen stimuli can be reliably decoded (along the diagonal) and hence evoke a non-sustained perceptual representation.

In terms of off-diagonal decoding, King and collaborators [24] previously reported temporal generalization for unseen stimuli. In their work, subjects had to compare the orientation of a masked Gabor to a subsequent probe following a brief delay period. Temporal generalization was observed for the presence versus absence of the target during the retention interval, even when subjects reported no subjective experience of the stimulus (although classification magnitude and duration was considerably reduced when compared with consciously seen trials). Such a finding implies that temporal generalization may be an imperfect index of conscious access, since it can occur – at least in part – for stimuli that were subjectively unseen, perhaps due to a short-term remanence and decay of neural codes that were activated subliminally. Our cross-phase generalization data from study three may be informative here, since they showed that training a classifier on task-relevant stimuli of phase three explained only the early portion of brain activity (∼180ms) when tested on task-irrelevant stimuli of phase one in subjects who were inattentionally blind to the stimuli. Note that a similar profile of late sustained activation for reported stimuli, but only early short-lived transient activity for unreported stimuli, has been previously reported in the attentional blink [59]. Such a pattern could, for example, indicate local recurrency without long-range recurrent processing for stimuli that remain unseen, and hence show that localised recurrent activity is sufficient for brief bouts of temporal generalization of unseen information at early timepoints. Yet there are also several key differences in the experimental design between studies that suggest temporal generalization may nevertheless remain uniquely associated with conscious processing in the specific experimental conditions examined here. Most notably, unseen trials in the work of King and colleagues [24] were report-based, where the unseen stimulus was always a task-relevant target. Comparatively, unseen trials in the datasets examined here always occurred under conditions of no-report, where stimuli were instead task-irrelevant non-targets. Since the distinction between report and no-report conditions is important in terms of the mental maintenance of perceptual representations, it stands to reason that it will also be important for whether classification of unseen stimuli can meta-stabilize and hence generalize across time (rather than show a serialized diagonal response). Indeed, in the work of King and colleagues [24], performance of classifiers trained on task-irrelevant sensory features of the target (e.g., spatial frequency) was at chance during the same delay period in which there was temporal generalization for the presence versus absence of the stimulus, providing some hint that unseen information that is task-irrelevant may not be afforded a capacity to stabilize across time. Evidently, more work is needed to understand whether and when temporal generalization occurs for task-irrelevant stimuli that remain unseen.

A similarly important question remains as to whether these findings generalise to other tasks and paradigms. Indeed, while an important strength of our findings is that they converged over multiple datasets, the fact that all datasets came from no-report inattentional blindness experiments calls to question whether these temporal generalization effects can be observed across any/all experimental settings where conscious content is manipulated, or whether they are instead unique to these experimental settings. So far, square, spreading, and/or thick diagonal patterns of generalization (see Figure 1) have also been found to be associated with conscious processing in other paradigms [e.g. 24-26, 28]. Yet, in a very recent set of results from a no-report masking experiment, temporal generalization was observed in the report condition while the no-report condition showed only diagonal decoding [16]. One possibility is that masking disrupted temporal generalization due to the suppression of informational persistence of the perceptual content in the underlying neural signal by the subsequent mask, which was always consciously perceived (but see [24]). Testing for the off-diagonal patterns observed here with different manipulations of awareness (e.g., stimulus degradation, attentional blink, binocular rivalry, etc.) will be especially important to ensure they are authentic signatures of conscious access independent of experimental design and may afford insight to better understand under what conditions temporal generalisation occurs for unconscious or subliminal stimuli. Relatedly, distinguishing diagonal from off-diagonal decoding is, at present, largely dependent on a qualitative judgment of the magnitude and spread of decoding, (i.e., to what extent a “thick diagonal” is observed). To more reliably assess for the presence of off-diagonal decoding—that is, when off-diagonal decoding should *not* be interpreted as temporal generalisation of decoding—future work could look to establish a quantitative metric to distinguish one from the other.

## Conclusion

The current set of results reveal a robust electrophysiological signature of conscious processing through temporal generalization of decoding in three distinct no-report inattentional blindness paradigms over four previously published datasets. These multivariate patterns show some consistency across stimulus types ranging from very basic shapes to more complex stimuli (faces, words) and are observable independent of report requirements and the evocation of a P3b. Our findings suggest a complex, dynamic neural mechanism of conscious access that operates within a 200-400 millisecond post-stimulus time window, thereby strongly constraining the timing of conscious processing. Future work ought to explore these temporal generalization patterns across different experimental paradigms, stimulus durations, stimulus categories, and potentially even clinical populations, to further elucidate the fundamental neural processes underlying conscious experience.

## Authorship contributions

B.T.H. and M.P conceived the project. B.T.H. performed all analyses and drafted the manuscript. M.P. provided datasets and supervision. M.P., S.D., and H.S. provided review, editing, and analytical expertise and feedback.

## Methods

### Participants

A total of 137 subjects from four studies were included in our sample, 75 (54.7%) of which were classified as “aware” and 62 (45.3%) as “inattentionally blind” to the unexpected critical stimuli in the first phase of the experiment. Thirty-two subjects (*M* age = 21, 19 females) were included from study one [3], 16 of which were classified as aware and 16 as inattentionally blind. Study two had a sample of 31 subjects (*M* age = 19.96, 13 females), 16 of which were classified as aware and 13 as inattentionally blind [32] (three subjects were excluded at the analysis stage due to insufficient trials). Study three’s sample of 30 subjects (*M* age = 21, 21 female) was made up of 15 aware and 15 inattentionally blind subjects [5]. Study four’s sample included 46 subjects (*M* age = 20.02, 26 female), 28 of which were classified as aware and 18 as inattentionally blind [6]. For all studies, subjects had normal or corrected-to-normal vision. Subject participation in all studies was voluntary and all were remunerated with either cash or course credit. All studies received approval from the relevant ethics IRBs.

### Procedure

Each study used the same three-phase macro set-up that involved two separate tasks. Subjects first ran through several practice trials (where no critical stimuli were presented) before proceeding to the main experiment. Blocks of trials lasted between 60 and 120 s. Subjects held their gaze at the centre of the screen throughout each experiment. In practice trials and in phase one and phase two, subjects had to detect with a button press a change in luminance of one of the discs (see Figure 2 and Stimulus Materials), while in phase three subjects had to attend and respond with a button press to a target pattern within the line segment arrays: the diamond pattern for study one, faces with a missing feature for study two, and animal words in study three. While sequence duration varied slightly between studies, the key stimuli were always presented for 300ms. In each experiment, the distractor task (disc detection) was used to induce inattentional blindness to the critical stimuli in phase one (in approximately half of the subjects), with the awareness assessment between phase one and two (see below) serving as both a retrospective examination of inattentional blindness and as a cue to induce awareness in all subjects in phase two. The task change from phase two to phase three altered the relevancy of the critical stimuli: critical stimuli (squares, faces, words/letters) were task-irrelevant in phase one and two – coinciding with the ‘no-report’ component of each study – and task-relevant in phase three. Targets for all phases / experiments occurred on 10% of trials, except for study three, where they occurred on 20% of trials.

Self- paced breaks were provided after every block. There was an extended break after phase one and two where subjects completed an awareness assessment probing their awareness of the unexpected critical stimuli retrospectively for the previous phase. Although there was some slight variation in the precise questions asked in the awareness assessments between studies, across all studies the assessment began by querying subjects with a forced choice item (yes / no) on whether they noticed any patterns in the background, followed by an open-ended item asking subjects to describe (or draw) what they saw (if anything). Following these items, example images of stimuli were shown, one by one with no time limit, and subjects had to rate their confidence in having seen the stimuli during the previous phase (1 = *very confident I did not see it*; 2 = *confident I did not see it*; 3 = *uncertain*; 4 = *confident I saw it*; 5 = *very confident I saw it*) and how frequently they had noticed them (1 = *never*; 2 = *rarely / less than 10 times*; 3 = *infrequently / 10–50 times*; 4 = *frequently / 50–100 times*; 5 = *very frequently / more than 100 times*). Five example stimuli were shown in a random order, including the critical stimulus, the stimulus that would eventually serve as a target in phase three, and three foil stimuli that never appeared. Subjects were classified as inattentionally blind if they answered “no” to the first question, left the second question blank (or sketched only randomly oriented lines in their drawing), AND rated their confidence in having seen the critical stimulus (when shown the example on the screen) as a 3 or less. Subjects were classified as aware if they answered “yes” to the first question OR sketched the critical stimulus in their drawing OR described it in writing when answering the second question OR gave a confidence rating above 3 after being shown the example critical stimulus on the screen. One subject from study three [6] who rated their confidence in having seen the foil stimuli above 3 on the scale was excluded from analysis. Overall, this approach to classifying subjects as inattentionally blind was designed to err on the conservative side to ensure a complete lack of awareness in the inattentionally blind group, while minimally risking some misclassification of inattentionally blind subjects into the aware group (rather than the other way around). In practice, it was fairly obvious which subjects were inattentionally blind, as most reported being surprised after seeing the critical stimuli in phase two, with many expressing shock and disbelief given how salient the stimuli were and how frequently they appeared. On several occasions, inattentionally blind subjects asked the experimenter questions such as: “did those faces really appear that many times in the first part of the study?” and “how could I have missed all of those shapes before?”.

### Stimulus Materials

Each study used broadly similar experimental presentation parameters where many line segments were displayed that could vary between several possible configurations. In each, line segments formed: (1) one stimulus type that served as the unexpected critical stimulus in phase one, the expected (but still task-irrelevant) critical stimulus in phase two, and the task-relevant non-target stimulus in phase three, (2) one stimulus type that was the task-relevant target in phase three, and (3) one stimulus type that served as the “control” stimulus throughout all phases (see Table 1). These three types of stimuli were always presented for 300ms, while an array of randomly oriented line segments was presented during the inter-stimulus intervals (600-800ms for studies one and three; 533-600ms for study two) (see Figure 2). Distractor stimuli were displayed either overlaying the line segments (study two/three) or peripheral to the line segments (study one), such that subjects had to covertly attend to distractor stimuli while maintaining fixation at the centre of the screen. Stimuli (including the distractors) were physically identical and presented for the same numbers of trials across all phases, with the only difference being the randomly shuffled order of presentation of the three stimulus types within each phase.

**Table 1.** Study Design Parameters for each Iteration of the No-Report Inattentional Blindness Experiment.

| Study | Dataset | Stimuli per phase | Trials per block | Blocks per phase | Phase duration | Distractor stimuli | Critical stimuli | Target phase one/two | Target phase three |
| --- | --- | --- | --- | --- | --- | --- | --- | --- | --- |
| One | One [3] | 600 | 60 | 10 | 10 min | Peripheral red ring with eight red discs | Square patterns (40%) | Red disc luminance decrease (10%) | Diamond array (10%) |
|  | Two [32] | 600 | 60 | 10 | 10 min | Peripheral red ring with eight red discs | Square patterns (40%) | Red disc luminance decrease (10%) | Diamond array (10%) |
| Two | Three [5] | 720 | 72 | 10 | 12 min | Three green rings with 12 green dots | Faces (30%) | Green dot luminance increase (20%) | Faces with missing feature (20%) |
| Three | Four [6] | 800 | 80 | 12 | 12 min | Three green rings with 12 green dots | Words / consonant strings (45%) | Green dot luminance increase (10%) | Animal words (10%) |

*Note.* Percentages in parentheses indicate the proportion of trials in which the stimulus appeared.

*Stimuli*. In study one [3], 400 (20 × 20) small, straight-line segments (each subtending approximately 0.34° for study one and 0.45° visual angle for study two) were displayed on a dark background (0.07 cd/m^2^) on an LCD monitor with a refresh rate of 60 Hz (study one viewing distance = 120cm, study two viewing distance = 57cm). The central grid of line segments varied between (1) a square pattern made from 44-line segments (3.5° x 3.5° visual angle), (2) a diamond array composed of 28-line segments (3.2 × 3.2° visual angle), and (3) a random configuration (each line segment orientation was assigned randomly from 15 steps in degrees of rotation). Squares were the critical stimuli and were presented on ∼40% of trials (240 presentations per phase), whereas diamonds were targets in phase three and were presented on ∼10% of trials (60 per phase). Random configurations were presented on the remaining ∼50% of trials (see Figure 2).

In study two and three [5, 6], the screen displayed hundreds of randomly oriented “squiggly” line segments presented on a white background on a 1920 ×1200-pixel display screen (Planar SA2311w23 LCD monitor) with a refresh rate of 60 Hz (study three viewing distance = 75cm, study four viewing distance = 72cm). In study three [5], line segments varied between: (1) a human face made from 36-line segments (5.4° visual angle) and presented on 30% of trials (216 presentations per phase), (2) face stimuli that had a feature missing (eye, nose, or mouth), presented on 20% of trials (144 per phase), and (3) randomly oriented control stimuli, made from the same line segments as faces but which were oriented such that no discernible pattern was present, presented on 50% of trials (360 per phase).

Intact faces were the critical stimuli and faces with missing features served as the targets in phase three. In study three [6], line segments formed (1) five-letter word stimuli made from 31 line segments centered inside of a larger 13 × 13 array of line segments (16.0° visual angle), presented on 22.5% of trials (180 presentations per phase), (2) five-letter animal words, presented on 10% of trials (80 per phase), (3) consonant strings, made up of any of the 21 consonants of the English alphabet, five letters long, presented on 22.5% of trials (180 per phase), and (4) randomly oriented arrays, which consisted of five ‘non-letter’ line segments made up of the same lines used to generate word/consonants, presented on 45% of trials (360 per phase). Words and consonant strings were the critical stimuli, while the animal word stimuli served as targets in phase three (see Figure 2). More details about the stimuli from each study can be found in the original reports [3, 5, 6].

*Distractor stimuli/task*. In study one, the line array grid was surrounded by a red ring of eight discs (each approx. 1° visual angle) that remained static for 600–800ms (during the inter-stimulus intervals) and then rotated either clockwise or counterclockwise by 15° to form one of two alternate positions for 300ms (during the stimulus intervals). On ten percent of the trials, the luminance of one of the eight discs decreased, which served as the target during phase one and phase two. In study two, three concentric green rings (largest 6.5°, middle 4.5°, smallest 2.5°), each containing four small green disks spaced equidistantly, overlaid the line segments; whereas in study three, three concentric green ellipses (height = 1.6°, 3.3°, 5.0°; width = 4.0°, 5.6°, 7.4°, respectively), each containing four evenly spaced green disks, overlaid the line segments. In both study two and three, disks were smallest on the innermost ring (0.32°), larger on the middle ring (0.41°), and largest on the outermost ring (0.52°).

Disks rotated continuously along the rings at a rate of 52°/s throughout each block of trials. The direction of rotation alternated between clockwise and counterclockwise across successive blocks (every 1 min), but never changed within a block. For study two, one of the green dots increased in luminance for 300ms on 20% of the trials, whereas this occurred on 10% of trials for study three.

### EEG

Dataset one [3] used a commercially available electrode cap with a 64-channel custom montage, where EEG was amplified by a battery-powered amplifier (SA Instrumentation, San Diego, CA) with a gain of 10,000 and band-pass filtered from 0.1 to 250 Hz. Electrode impedances were kept below 5 kΩ. Dataset two [32] used a standard 10-20 64-channel montage digitized using the BioSemi Active Two system and recorded at a sample rate of 1024 Hz. Dataset three [5] and four [6] used an EASYCAP custom 96-channel electrode cap with equidistantly spaced Ag/AgCl electrodes. EEG signals were recorded with three 32- channel amplifiers (Brain AmpStandard; BrainProducts), band-pass filtered online from 0.1 to 150Hz, and digitized at 500 Hz. Electrode impedances were kept below 5kΩ.

All pre-processing was completed using custom Matlab scripts written using EEGlab and MVPAlight [60, 61]. Datasets were re-referenced to the average of all scalp channels, down sampled to 250Hz, epoched to 1200ms time windows time locked to -200ms pre-stimulus to +1000ms post-stimulus onset (critical stimulus, random lines), and baseline corrected using a -100 to 0ms baseline. Note that trials time-locked to targets of phase three were *not* used in any dataset; only trials with the presentation of square patterns (study one), intact faces (study two), and words and consonant strings (study three) were used. In other words, trials with the presentation of diamonds (study one), missing feature faces (study two), and animal words (study three) were excluded from all analyses (due to the small numbers of trials). No other pre-processing was performed, other than a 50hz notch filter applied to the dataset of study two [32], since this was the only dataset collected in Australia (line noise: 50hz) and not within an electrically shielded, sound attenuated booth. Trials were removed across all studies that included a button-press response and/or target disc, leading to the following average number of trials per condition, study one (shapes): 185.5 (phase one), 187.1 (phase two), 197.2 (phase three); study two (faces): 129.9 (phase one), 131 (phase two), 111.6 (phase three); and study three (words/letters): 262.3 (phase one), 265.7 (phase two), 262 (phase three).

Decoding was performed using the MVPA light toolbox in Matlab [61]. Datasets were reformatted from a channel by time by trial structure, to a trial by channel by time structure, and labelled according to their respective conditions (critical stimulus, control). As part of MVPAlight’s built-in preprocessing parameters, we applied *under sampling* – where trials are removed from the condition with a higher number of samples (which here was always the control condition) to equate the number of samples between conditions, *z-scoring* – where data are demeaned and rescaled, and *average sampling* – where, to improve signal-to-noise ratio, trials from the same condition are divided into groups and each group is replaced with the group average. We used default parameters for each which, for average sampling, meant that trials were divided into five groups and that each ‘pseudo average’ was composed of approximately 4 trials on average.

For our classifier, we used a linear support vector machine with five iterations of a ten-fold cross validation procedure (five-fold cross validation was used for a few subjects, *n* = 7, due to insufficient trials for reliable folds with ten-fold cross validation) toward classification of critical stimuli from their controls. We implemented temporal generalization (where each classifier is trained on a given time point and tested on every time point) and searchlight analysis (where electrode sensor, rather than time, is treated as the dimension over which the classifier is trained/tested). Temporal generalization reflects the degree to which the classifier *generalises* across time, whereas searchlight indicates classification performance for different scalp locations for a given time window. Specifically, classification is repeated for each electrode separately, thereby obtaining a classification performance value for each electrode, which is then plotted to show the performance as a scalp topography (where neighbours were set based on a standard 64/96 channel layout).

We inspected off-diagonal decoding performance of classifiers trained from 200- 800ms in 100ms intervals (200ms, 300ms, 400ms, 500ms, 600ms, 700ms, 800ms) in order to gauge the evolution of temporal generalization over time. Our searchlight analysis complemented this by inspecting the scalp distribution of classification performance over four corresponding time windows (200-300ms, 300-400ms, 400-500ms, 500-600ms). Note that because dataset one [3] and two [32] used different electrode montages, for simplicity we present searchlight analysis results from study one only.

Model performance was assessed using area under the curve (AUC), which can be interpreted similar to accuracy but is robust to differences in offset and scaling across datasets. For statistical analyses of temporal generalization, performance was assessed using a two-tailed non-parametric Wilcoxon rank test performed at every time point within an epoch, with a cluster-based permutation for correction for multiple comparisons (1,000 permutations).

## Data Availability

The data that support the findings of this study are available from the corresponding author upon reasonable request.

